# High salt intake activates the hypothalamic-pituitary-adrenal axis, amplifies the stress response, and alters tissue glucocorticoid exposure in mice

**DOI:** 10.1101/2022.03.04.481654

**Authors:** Hannah M Costello, Georgios Krilis, Celine Grenier, David Severs, Jessica R. Ivy, Mark Nixon, Megan C Holmes, Dawn E.W. Livingstone, Ewout J Hoorn, Neeraj Dhaun, Matthew A Bailey

## Abstract

High salt intake is common and contributes to poor cardiovascular health. Sustained cortisol excess also induces an adverse cardiovascular profile. Urinary cortisol excretion positively correlates with urinary sodium excretion. We hypothesised that this was due to hypothalamic-pituitary-adrenal axis activation by high salt intake.

In male C57BL6/J mice, 2 weeks of high salt intake increased *Crh* and *Pomc* mRNA abundance in the hypothalamus and anterior pituitary, respectively and caused a sustained rise in plasma corticosterone. Plasma copeptin and anterior pituitary V1b receptor mRNA expression was elevated, which may contribute to basal HPA axis activation. Additionally, high salt intake amplified glucocorticoid response to restraint stress, indicative of enhanced HPA axis sensitivity. In the periphery, high salt intake reduced the binding capacity of corticosteroid-binding globulin, enhancing glucocorticoid bioavailability. Within several tissues, the expression of glucocorticoid-regenerating enzyme, 11β-hydroxysteroid dehydrogenase type 1, was increased and the glucocorticoid receptor downregulated. Overall, high salt intake increased glucocorticoid exposure in the hippocampus, anterior pituitary and liver.

Chronic high salt intake amplifies basal and stress-induced glucocorticoid levels and resets glucocorticoid biology centrally, peripherally and within cells. This shows direct connectivity between salt homeostasis and HPA axis function. The cumulative effect is likely maladaptive and may contribute to the long-term health consequences of a high salt diet.

## INTRODUCTION

The American Heart Association advocates a daily limit of 1,500 mg Na (3.75 g salt) for most individuals, particularly those with hypertension^1^. Daily salt intake in the USA, and most other countries, usually exceeds this threshold^2^. In humans and experimental animals high salt intake induces abnormal physiological function of many organ systems and causes a range of poor health outcomes^3^. This damaging effect of salt is most clearly seen in “salt-sensitive” experimental models, ie those which display an exaggerated hypertensive response to high salt intake. Salt-sensitivity is also found in ~30% of healthy humans and independently increases cardiovascular risk^4^ and mortality risk^5^. Nevertheless, even in those people categorised as salt-resistant, high salt intake modifies organ function and accrual of incremental deficits over time will cause disease. Thus habitual dietary salt excess causes cardiovascular disease and may contribute to the development of autoimmunity^6^, some cancers^7^ and cognitive impairment^8^. High dietary salt clearly exerts a significant global health burden. Interventions to reduce intake improve outcomes^9^. However, such reductions are difficult to achieve and sustain in the general population. Understanding how cell and organ physiology responds to chronic high salt intake offers an additional route to improve health across an array of diseases.

This study focuses on glucocorticoids (cortisol in humans, corticosterone in rodents), powerful hormones which underpin many important cardiovascular, cognitive, immune and metabolic cell functions. Glucocorticoids are not normally considered key regulators of salt balance or extracellular fluid volume homeostasis. Nevertheless, observational studies in humans show a positive correlation between urinary free cortisol excretion and 24-hour sodium excretion, taken to reflect salt intake^10, 11^. A small number of controlled sodium intake studies, typically lasting ~7 days, have examined the relationship between salt intake and urinary excretion of glucocorticoids. Ranging from 8 to >600 subjects, these studies consistently report a direct relationship between dietary salt intake and urine cortisol excretion^12–17^. A long-term balance study also found a positive relationship between salt intake and urinary cortisol excretion in healthy men enrolled in the MARS-500 spaceflight simulation programme^18^.

Although suggestive, these studies do assess the effect of high salt intake on circulating cortisol or glucocorticoid signalling within tissues. Here, we used C57BL6/J mice to test the hypothesis that sustained high salt intake would activate the hypothalamic-pituitary-adrenal (HPA) axis. We find that high salt intake caused multi-level disturbances in glucocorticoid biology that combine to increase plasma levels of hormone. This circulating and tissue-level excess increased glucocorticoid exposure in the liver and areas of the brain, which may contribute the adverse effects of sustained high salt intake.

## METHODS

Experiments were performed between September 2018 and September 2021. Adult male C57BL6/J mice were commercially sourced (Charles River, UK) at age 10-12 weeks and maintained under controlled conditions of temperature (24±1°C), humidity (50±10% humidity) and light (lights on 7am to 7pm local time). Mice had *ad libitum* access to water and commercial rodent chow. The control diet contained 0.3% Na and 0.7% K by weight (RM1, SDS Diets, UK); the high salt diet contained 3% Na and 0.6% K by weight (RM 3% Na^+^ SY, SDS Diets, UK). Mice were randomized into treatment groups and experiments performed with a single blind to group allocations.

The first experiment exposed mice to high salt intake for 2 weeks and assessed plasma corticosterone at diurnal peak and nadir. In other experiments, corticosterone was measured at the diurnal peak in mice fed either high salt or control diet for up to 8 weeks. For these experiments, mice were single-housed with access to bedding and enrichment. At the end of the experiments, animals were humanely killed for tissue collection. All experiments were performed in accordance with the UK Animals (Scientific procedures) Act under a UK Home Office Project License to the lead investigator (MAB) following ethical review by the University.

### Restraint stress testing

n=20 mice were used. Each animal was removed from their home cage between 7.15-8am with minimal disruption and 30μL blood collected within 1 min by tail venesection to give baseline corticosterone. Each mouse was then held in a Plexiglas restraint tube for 15 min, another blood sample taken for the peak corticosterone response, and then returned to the home cage. We were permitted to one further blood sample from each mouse and we measure plasma corticosterone at either 30-60- or 90-min post-restraint stress, giving n=6/7 at each time point. Statistical analysis of the post-restraint recovery phase was performed independently of the initial stress response.

### Corticosterone measurement

Corticosterone was extracted from tail venesection samples and measured by commercial ELISA kit (ADI-900-097; Enzo Life Sciences, UK), with a lower limit of detection of 27pg/ml.

### Aldosterone & copeptin measurement

Following decapitation (between 0715 and 0800 local time), trunk blood was collected on ice, and separated by centrifugation and stored at −20°C. After a single thaw of all samples, aldosterone (ADI-900-173; Enzo Life Sciences, UK; sensitivity of detection 4.7pg/ml) and copeptin (CEA365Mu; Cloud-Clone Corp., USA; lower limit of detection 9pg/ml) were measured by commercial ELISA.

### CBG binding capacity

CBG binding capacity was measured using a saturation binding assay, as described^19^. Terminal plasma samples (diluted 1:100) from mice (n=7/diet) were stripped of endogenous steroids using dextran-coated charcoal. CBG specific binding was calculated by subtracting non-specific binding from total binding. To measure non-specific binding, samples were co-incubated with saturating concentrations of unlabelled corticosterone. Unbound [1,2,6,7^-3^H]corticosterone was removed by further incubation with dextran-coated charcoal. The remaining radioactively labelled corticosterone bound to CBG was quantified by scintillation spectrophotometry. The maximal binding capacity (Bmax), measured in disintegrations per minute, was estimated using non-linear regression.

### RNA isolation and quantitative PCR

Tissues were snap frozen in dry ice and stored at −80°C. At use, tissues were homogenized from frozen and RNA isolated using RNeasy kits (Qiagen, US). RNA was quantified by the NanoDrop-1000 (Thermo Fisher Scientific, UK) and 500ng cDNA was generated using high-capacity RNA-to-cDNA kit in 10μL reactions (Applied Biosystems, UK). The mRNA expression was measured by quantitative RT-PCR using the Universal Probe Library (Roche, UK). Primer sequences and probe numbers are given in **Supplemental table 1**. Triplicates of each sample and standard curve were run on the LightCycler480 (Roche, UK). The following conditions were used: 95°C for 5 min as preincubation, 95°C for 10 sec and then 60°C for 30 sec for a 50-cycle program, followed by a cooling period of 30 sec at 40°C. The results were normalized to the mean concentration of reference genes (*Actb* and *Tbp* for adrenal, hippocampus and anterior pituitary; *Actb* and *Hprt* for liver; *Actb* and *Rn18s* for kidney cortex/medulla and aorta; *Actb* and *Gapdh* for heart; *Gapdh* and *Tbp* for hypothalamus). There was no effect of diet on the mRNA abundance of the reference genes.

### Quantification and statistical analysis

Group size estimates were based on pilot experiments demonstrating an increase in peak plasma corticosterone after 2 weeks of high salt intake. Power analysis was performed using G*Power v3.19 Software^20^, worked to a main effect alpha level of *p* <0.05 and 90% power. No animals were excluded from study but some samples subsequently failed quality control in assay (eg for RNA quality/abundance) and retrospective power analysis on datasets was therefore performed. In final data sets, individual points are shown along with group mean±SD. Statistical analysis was performed using GraphPad Prism v8.4 Software. The distribution of data sets was assessed using the Shapiro-Wilk normality test. Normally distributed datasets were analysed by *t*-test, one-way ANOVA or two-way ANOVA, with or without repeated measures, as stated in the figure legends. For ANOVA, Holm-Sidak post-tests were used for planned comparisons only when the main effect *p* <0.05. For data that did not pass normality testing, non-parametric analysis was carried out using Wilcoxon test. The sample number (*n*) and statistical analysis details used are stated in the figure legends; absolute *p* values for planned comparisons are reported.

## RESULTS

### High salt activates the HPA axis to increase corticosterone synthesis

The hypothesis that high salt intake activated the HPA axis was initially tested in n=8 mice, sampling blood at 7am and 7pm to capture the diurnal nadir and peak of corticosterone (Figure 1A). Measurements were made in each mouse on control diet (0.3% Na) and again after 2 weeks of high salt (3% Na). High salt intake significantly increased peak plasma corticosterone, without affecting the nadir (**Figure 1A**). We next examined the time course, measuring peak corticosterone in separate groups of mice maintained on high salt or control diets for 3 days, 1-, 3-, and 8-weeks (**Figure 1B**). The effect of salt intake was biphasic: at 1 week, peak corticosterone was supressed, after which stimulatory effect of high salt intake was sustained.

**Figure 1:**
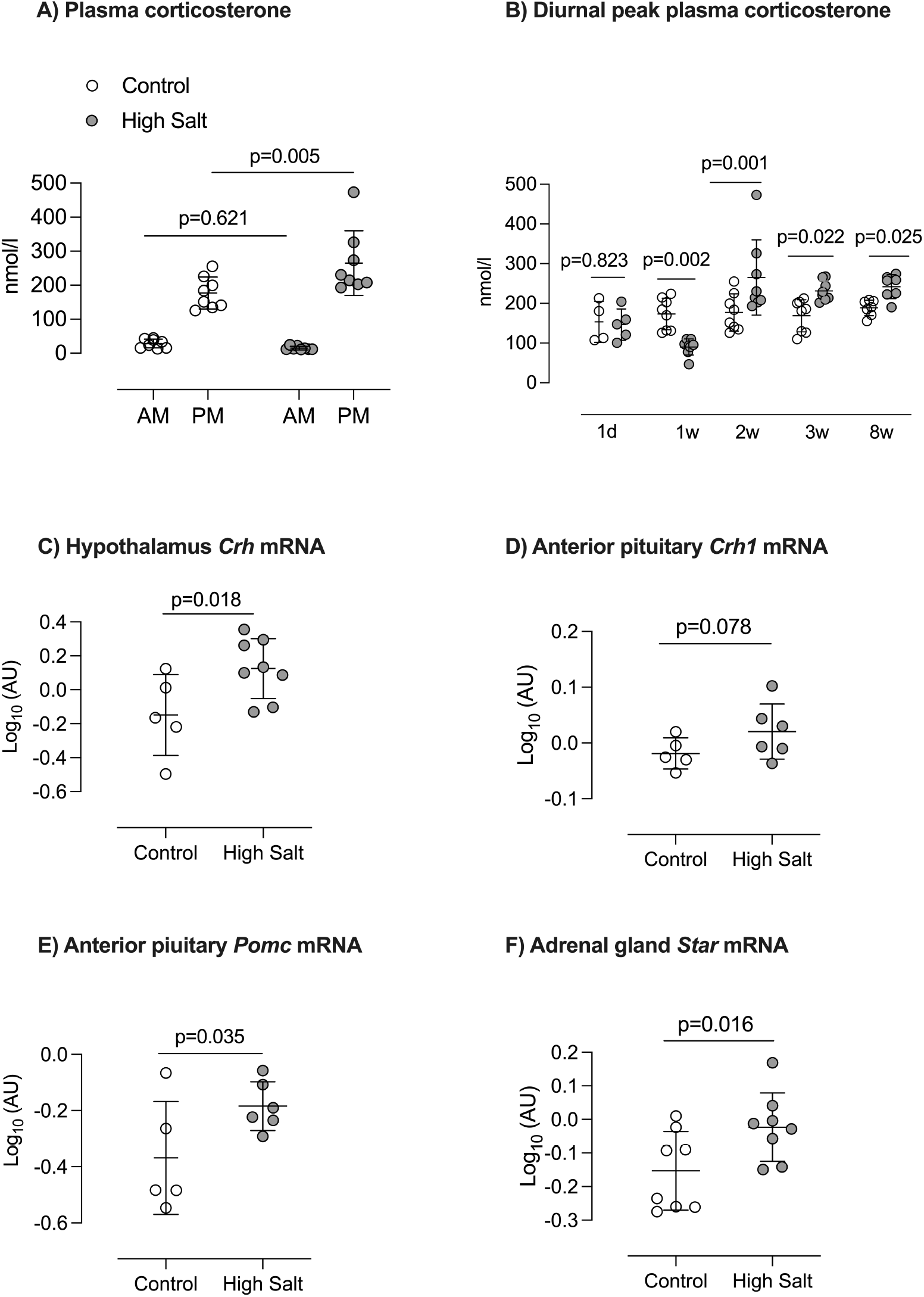
Activation of HPA axis after high salt intake. **A)** Plasma corticosterone measured at 7am (diurnal nadir) and 7pm (diurnal peak) local time in male C57BL6 mice (n=8) first on a 0.3% Na diet (Control, open circles) and again after 14 days on 3% Na diet (High Salt; grey circles). Individual values are shown with group mean±SD; statistical comparison was two-way repeated measures ANOVA for the main effects of time of day (p<0.0001) and diet (p=0.071) and the interaction (p=0.016); p-values for planned comparisons (Holm-Sidak) are given. **B)** The effect of salt diet on diurnal peak plasma corticosterone was measured in separate groups of mice at 3 days, 1, 2, 3 and 8 weeks of either control or high salt intake. Individual values are shown with group mean±SD; statistical comparison was two-way ANOVA for the main effects of “*diet*” (p=0.026) and “*duration*” (p<0.0001) and the interaction (p<0.0001); p-values for planned comparisons (Holm-Sidak) are shown. To test the hypothesis that the HPA axis was activated, mRNA abundance was measured for **C)** *Crh* (corticosterone releasing hormone); **D)** *Crh1* (CRH receptor); **E)** *Pomc* (pro-opiomelanocortin); and **F)** *Star* (Steroidogenic Acute Regulatory protein) in tissue taken from different C57BL6/J mice fed either 0.3% Na (open circles) or 3%Na (grey circles) for 14-days prior to cull. Individual values are shown with group mean±SD; statistical comparison was by unpaired t-test with one-tailed p-values as indicated.

Using tissue taken from the 2-week high salt group, we demonstrated that hypothalamic *Crh* expression (encoding corticotropin releasing hormone; CRH) was elevated (**Figure 1**). The anterior pituitary CRH receptor, *Crhr1*, was not transcriptionally increased by high salt intake but pro-opiomelanocortin, the precursor of ACTH, was. Adrenal gland weight was not changed by high salt diet, nor was the expression of the ACTH receptor (**Supplemental Figure 1**). mRNA abundance for *Star* was increased (**Figure 1**). This encodes Steroidogenic Acute Regulatory protein, which is rate-limiting for adrenal steroid biosynthesis. Nevertheless, the stimulatory action was limited to glucocorticoid and there was broad suppression of the aldosterone system by high salt intake: plasma aldosterone was reduced, as was mRNA for hepatic angiotensinogen (*Agt*) and adrenal gland aldosterone synthase (*Cyp11b2*); the renal expression of the mineralocorticoid receptor was significantly lower in mice on a high salt diet (**Supplemental Figure 2**).

In a different group of mice (n=20), we assessed the effect of high salt intake on the HPAA response *to* a 15-minute restraint stress. On control diet, restraint stress increased plasma corticosterone in all mice, with a group average of ~90 nmol/L (**Figure 2A**). After 2-weeks of high salt intake, the peak stress response was amplified in 16 mice (**Figure 2B**), and the group approximately doubled. Protocol welfare restrictions restricted us to only one additional blood sample per mouse, drawn by tail venesection at either 30-, 60- or 90-minutes post-restraint (**Figure 2C**). Assessed by ANOVA at group level, the main effects of time (p<0.0001) and diet (p=0.007) were significant, but the interaction was not (p=0.344) and there were no significant differences in the planned *post-hoc* comparisons, indicating that the rate of recovery from stress was not affected by salt intake.

**Figure 2:**
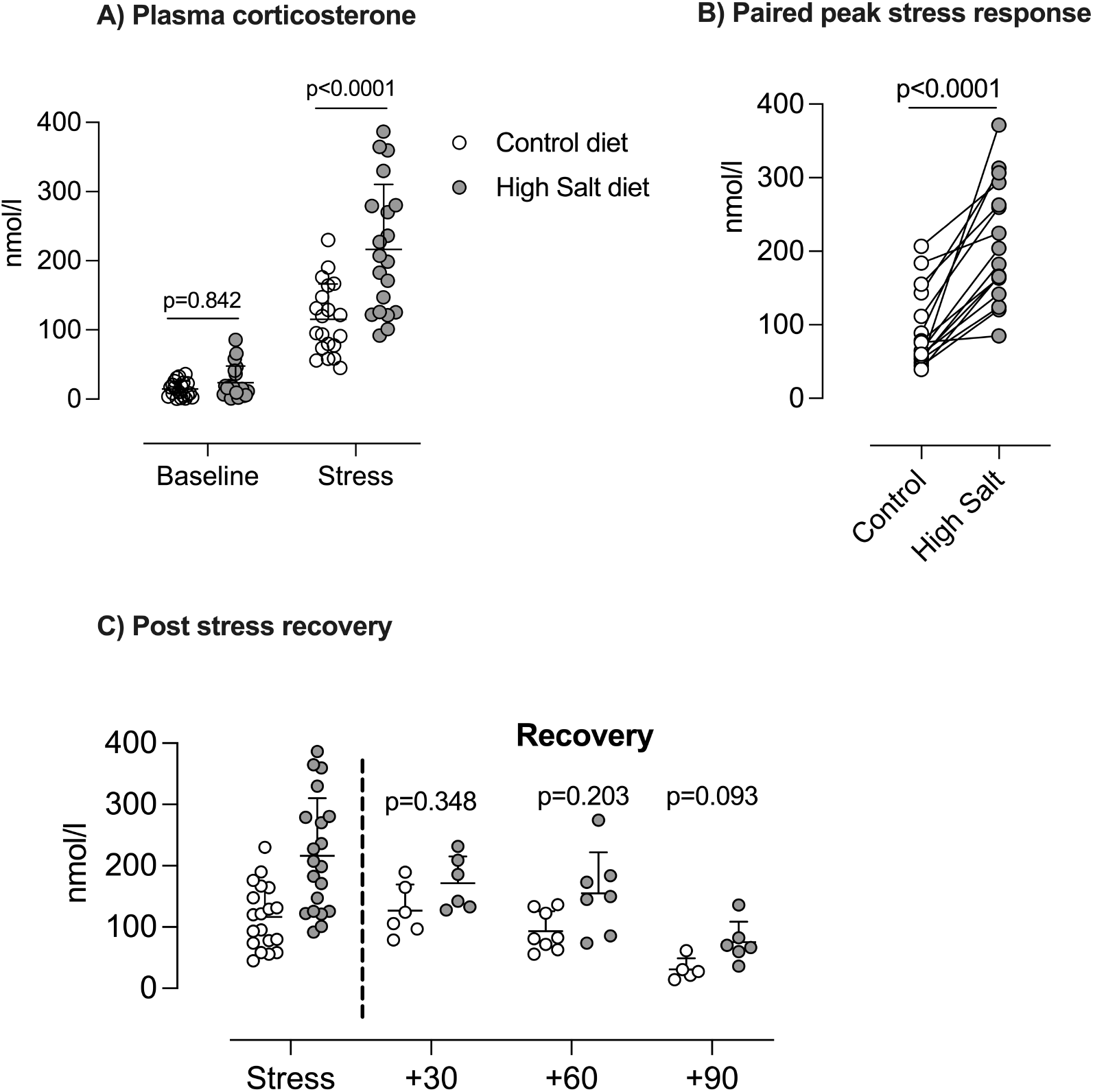
High salt intake amplifies the stress response. **A)** Plasma corticosterone was measured in male C57BL6/J mice (n=20) fed 0.3% Na diet (Control; open circles) at 9am (baseline) and again after 15 minutes of tube restraint (Stress). Measurements were repeated after 2 weeks on a 3% Na diet (High Salt; grey circles). Analysis was two-way ANOVA with repeated measures for the main effects of “*stress-response*” (p<0.0001) and “*diet*” (p=0.0006) and the interaction (p=0.0014); p-values for planned comparisons (Holm-Sidak) are given. **B)** The paired peak stress response in each mouse, analysed by pair-t-test. **C)** The corticosterone recovery from stress was measured at either 30, 60 or 90 minutes following release from the restraint tube. Analysis was two-way ANOVA with repeated measures for the main effects of “*time*” and “*diet*” and the interaction; p-values for planned comparisons (Holm-Sidak) are given. Individual measurements are shown with group mean±SD.

After 14 days of high salt intake plasma osmolarity was increased, as was copeptin (**Figure 3**), a stable surrogate for vasopressin secretion^21^. Vasopressin, a neuropeptide, acts centrally *via* V1b receptors to stimulate ACTH release and peripherally *via* V2 receptors to stimulate renal sodium and water reabsorption. In these mice, both anterior pituitary V1b receptor and renal V2 receptor mRNA abundance was higher than in those on control diet. There was no difference in body weight between groups at the start or end of the experiment; high salt diet induced significant polydipsia and polyuria.

**Figure 3:**
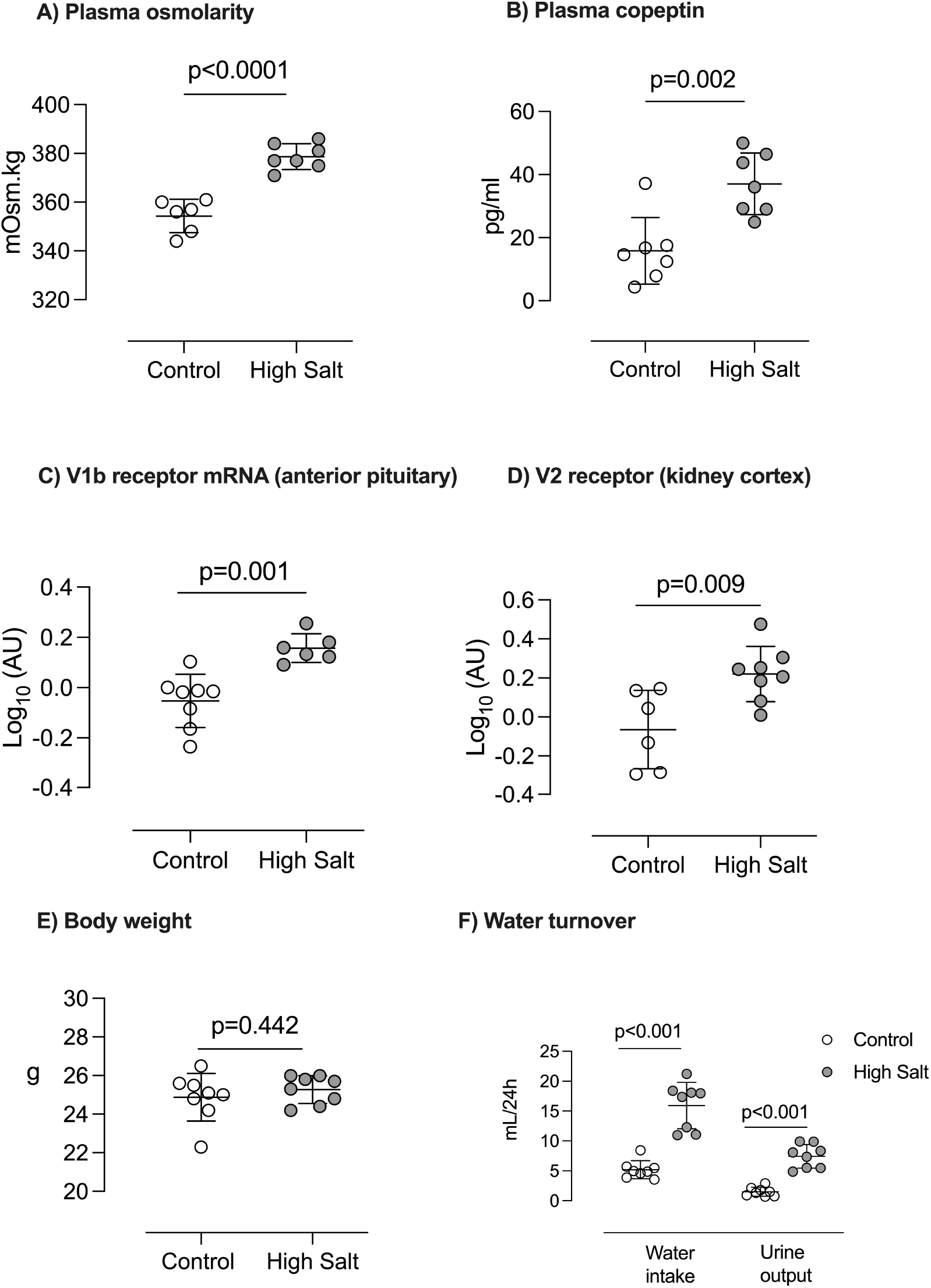
Effect of high salt intake on fluid volume markers. A) plasma osmolarity; B) plasma copeptin; C) anterior pituitary V1B receptor mRNA abundance; D) renal V2 receptor mRNA abundance; E) body weight and F) water turnover in male C57BL6/J mice fed either 0.3% (Control; open circles) or 3% (High Salt; grey circles) diet for 14 days. Individual measurements are shown with group mean±SD; statistical comparisons were made with Student’s unpaired t-test and two-tailed p-values are given.

### High salt reduces Corticosterone Binding Globulin expression and capacity

High salt diet activates the HPA axis to increase circulating glucocorticoid at rest and under stress. However, most glucocorticoid circulates bound to corticosterone-binding globulin (CBG), which can buffer against increased hormone production. After 2 weeks of high salt intake, CBG binding capacity was reduced and hepatic expression of the encoding gene, *SerpinA6*, was lower by >50% (**Figure 4**). *SerpinA6* expression was also significantly reduced in mice after 1 week of high salt intake (**Supplemental Figure 3**); binding capacity was not altered at this time point.

**Figure 4:**
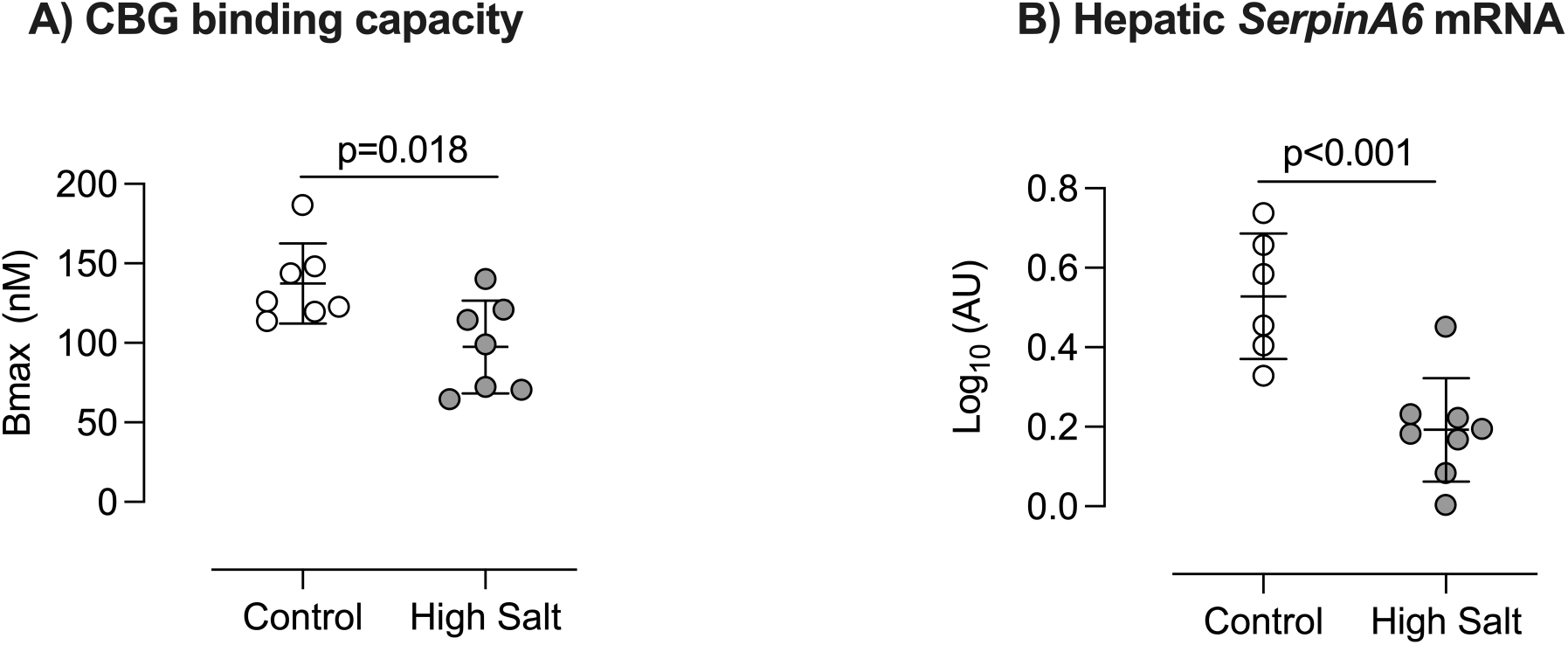
Effect of high salt intake on Corticosterone Binding Globulin. Male mice were fed either 0.3% (Control; open circles) or 3% (High Salt; grey circles), diet for 14 days and blood taken for measurement of A) Corticosterone Binding Globulin binding capacity. B) mRNA abundance of *SerpinA6*, encoding CBG, was measured using liver cDNA from another experimental group. Expression was normalized to that of reference genes *Gapdh* and *Tbp* and log-transformed. Individual measurements are shown with group mean±SD; statistical comparisons were made with Student’s unpaired t-test and two-tailed p-values are given.

### High salt alters glucocorticoid receptor signalling in tissues

High salt diet down regulated glucocorticoid receptor mRNA in bulk homogenates of most of the tissues examined (**Figure 5, Supplemental Figures 4-7**). In some tissues, the enzyme 11βHSD1, which regenerates active glucocorticoid in cells and amplifies glucocorticoid signalling^22^, was upregulated at mRNA level. Tissue glucocorticoid exposure, reported by *Fkbp5* expression^23^ was increased in the hippocampus, anterior pituitary and liver, downregulated in the kidney and not significantly changed in the heart, aorta, adrenal and white adipose tissue after two weeks of high salt intake.

**Figure 5:**
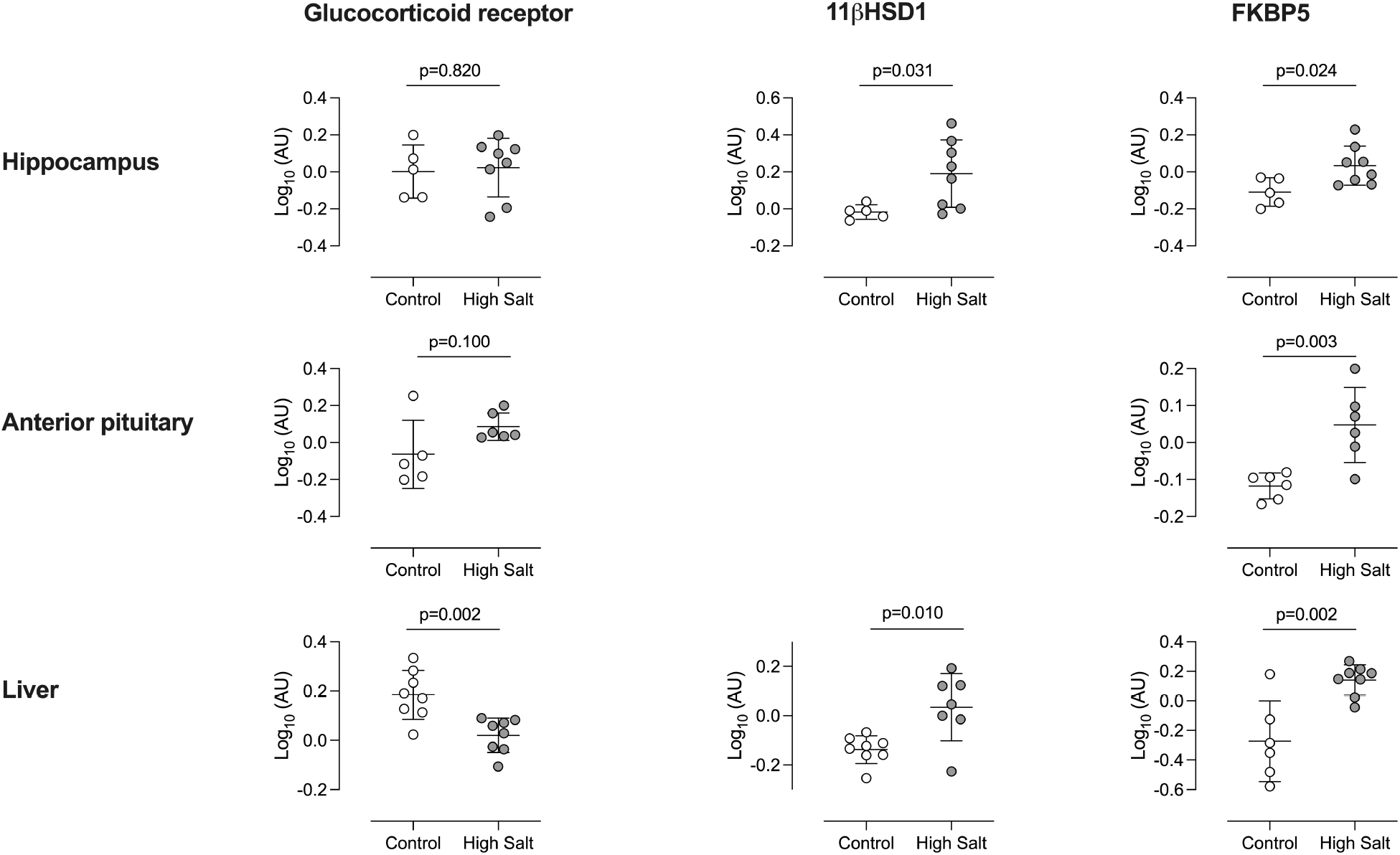
Effect of high salt intake on organ glucocorticoid exposure. Male mice were fed either 0.3% (Control; open circles; n=8) or 3% (High Salt; grey circles; n=8), diet for 14 days and tissues taken at cull. mRNA abundance of glucocorticoid receptor (Nr3c1) and FK506 binding protein 5 (*Fkbp5*) was measured in A) hippocampus; B) anterior pituitary and C) liver. 11β hydroxysteroid dehydrogenase type 1 (*hsd11b1*) was also measured in hippocampus and liver. Values were normalized to the expression of reference genes *Gapdh* and *Tbp* and log-transformed. Individual measurements for samples that passed QC are shown, along with group mean±SD. Statistical comparisons were made with Student’s unpaired t-test and two-tailed p-values are given.

## DISCUSSION

Sustained glucocorticoid excess causes multi-morbidity and poor cardiovascular outcomes^24, 25^. The major finding of our study is that sustained high dietary salt intake in mice induced multiple abnormalities in glucocorticoid biology. The HPA axis was activated and the forward response to environmental stress amplified, whereas the capacity to buffer blood corticosterone through binding to CBG was reduced. This study provides an explanation of the positive correlation between salt intake and urinary cortisol excretion consistently observed in humas^12–18^, and has two important implications. First, glucocorticoid excess may contribute to the long-term health consequences of high salt intake; second, dietary salt intake becomes an important consideration when diagnosing and managing preexisting hypercortisolism.

### HPA axis activation

We show functional evidence of a novel, direct connection between dietary salt intake and HPA axis activation. The effect was biphasic: diurnal peak plasma corticosterone was supressed at 1 week and thereafter significantly enhanced. In humans, 1 week of high salt also reduces plasma cortisol and this may reflect an initial phase of enhanced urinary elimination^26^. Longer-term, however, our data suggest that high salt intake increases corticosterone production: peak plasma levels were elevated, coupled with upregulation of hypothalamic CRH. Moreover, there was no downregulation of CRH receptor expression in the anterior pituitary, which normal occurs with sustained increases in circulating glucocorticoid^27, 28^. Indeed, the increase in anterior pituitary *Pomc* mRNA is indicative of enhanced ACTH production/release with sustained high salt intake. The drivers are not known but vasopressin, a co-regulator of ACTH secretion^29^, may contribute to HPA axis resetting. High salt intake increased serum copeptin, which quantitatively reports hypothalamic synthesis/release of vasopressin^21^, and also upregulated mRNA for the vasopressin V1b receptor in the anterior pituitary. Although *V1br* mRNA abundance does not fully parallel receptor number^30^, *V1br* knockout mice have reduced basal corticosterone and an attenuated response to acute stress^31^. Moreover, glucocorticoids directly increase mRNA *V1br* mRNA abundance and enhance receptor coupling to phospholipase C^32^.

Vasopressin synthesis is stimulated by hypertonicity and hypovolemia. After two weeks of high salt plasma osmolarity was significantly increased. It is difficult to understand. High salt rapidly induces polydipsia and polyuria but body weight is unchanged and increased water turnover does not seem to affect total fluid volume. However, high salt also increased renal aquaporin 2 expression, suggesting an intact renal response to perceived dehydration. High salt induces a complex osmotic response plasma response The fluid/volume response to high salt is complex^33^ and sustained exposure increases plasma sodium concentration in humans.^34^ The effect size is small (~3mmol/l) but modest changes are sufficient to activate hypertonic signalling in tissues.^35^

It is tempting to speculate that alterations in sodium and fluid-volume plasma stimulate vasopressin synthesis, synergising with CRH^36^ to activate the HPA axis. However, ACTH secretion is normally regulated by vasopressin derived from parvocellular neurones released into the pituitary portal plexus. Osmotic stimuli, in contrast, act at magnocellular neurones to promote vasopressin synthesis and released into the systemic circulation. Magnocellular vasopressin can induce ACTH production^37^ but chronic osmotic stress typically <u>inhibits</u> ACTH release^38^ and also impairs the pituitary responsiveness to novel stress. Our data point to the opposite and it may be that elevated serum copeptin reflects a stress response, rather than a hyperosmotic response. Indeed, serum copeptin is a sensitive marker of moderate psychological stress in humans^36^ and salt-sensitive individuals display an exaggerated cortisol response to acute mental stress^39^. Amplification of the stress response is a novel finding from this study, consistent with our previous report of sympathetic nervous system activation by high salt intake^40^. Other studies in male C57BL6 mice show that high salt intake promotes active coping strategies in response to a swim stress^41^. Rodent-based research in this area is not extensive, yet points to natural consumption of a high salt diet as an important behavioural modifier^42^.

### Peripheral glucocorticoid homeostasis

After two weeks of high salt feeding, CBG binding capacity, which determines circulating free glucocorticoid and delivery to tissues^43^, was reduced. This most likely reflects a reduction in total CBG protein since hepatic *Serpina6* mRNA abundance was also supressed by high salt feeding. Glucocorticoid excess suppresses CBG production by the liver^43^ but we do not consider this to be causal here because downregulation occurred at 1 week when peak plasma corticosterone was supressed. Reduced CBG expression increases the amount of free corticosterone in circulation and increased glucocorticoid signalling might therefore be anticipated. However, CBG also mediates tissue delivery and CBG-deficient mice are hyporesponsive to corticosterone^44^. Our data do not indicate a consistently reduced glucocorticoid response. Indeed, elevated *Fkbp5* mRNA suggests glucocorticoid exposure was enhanced in liver, anterior pituitary and hippocampus. The overall picture is complex: systemic changes to production and binding were accompanied by downregulation of glucocorticoid receptor expression and, in some tissues, indication of enhanced intracellular glucocorticoid regeneration *via* 11βHSD1. Cellular level changes in glucocorticoid regeneration can be physiologically significant, particularly when coupled with systemic glucocorticoid excess^45^. For example, transgenic 11βHSD1 overexpression in liver^46^ or adipose^47^ cause metabolic abnormalities and salt-sensitive hypertension and overexpression in the hippocampus accelerates age-related cognitive decline^48^.

### What purpose does HPA activation serve?

A long-term balance study in humans identified a positive correlation between the rhythmicity of urinary cortisol and salt excretion at constant salt intake. It was proposed that cortisol stimulated renal salt excretion^18^ and one interpretation is that our study shows a coordinated, whole-body response through which glucocorticoids contribute to salt balance during the sustained challenge of high salt intake. Certainly, synthetic glucocorticoids can induce negative sodium balance^49^, largely attributable to hemodynamic effects^50^. However, glucocorticoid receptor activation stimulates renal sodium transport^51, 52^ and glucocorticoid excess typically promotes sodium retention and salt-sensitive BP abnormalities^53–55^. Moreover, high salt initially *reduced* circulating corticosterone. The glucocorticoid excess induced by persistent high salt intake does not fit to a rapid, homeostatic response to excrete salt and is more likely to be a maladaptive response to complex fluid/volume changes and/or perceived stress.

### Perspectives

Observational studies report a positive association between sodium intake and cortisol excretion in woman and men^10, 11^. We find evidence pointing to accumulated glucocorticoid excess with sustained high salt intake in healthy C57BL6 mice. Importantly, studies in humans^26^ and mice^56^ suggest that this relationship may be exaggerated in salt-sensitive individuals. Our study is limited to a description of a phenomenon and has not explored how HPA activation influences organ function when salt intake is high. The snap-shot view at 2 weeks indicates that the degradation of regulatory mechanisms in brain and blood is accompanied remodelling of cellular glucocorticoid signalling across many organs. We speculate that HPA activation will induce diverse changes across physiological systems, which may contribute to poor cardiovascular^3^, metabolic^10^,^57^ and cognitive health when salt intake is high.^58 50^

## Funding Sources

Research funding for this study came from The British Heart Foundation (FS/16/54/32730; PG/16/98/32568) and Kidney Research UK (IN001/2017; INT001/2018). ND is supported by a Chief Scientist Office Senior Clinical Research Fellowship (SCAF/19/02).

## Disclosures

None

## Notes

### Competing Interest Statement

The authors have declared no competing interest.

