## Supplemental Figures and Table for "High salt intake activates the hypothalamic-pituitary-adrenal axis, amplifies the stress response, and alters tissue glucocorticoid exposure in mice"

**Data Supplement for:**

**Table S1: List of Primers**

| **Gene** | Protein | Forward primer | Reverse Primer | UPL probe no. |
|---|---|---|---|---|
| *Actb* | βactin | ctaaggccaaccgtgaaaag | accagaggcatacagggaca | 64 |
| *Gapdh* | GAPDH | gggttcctataaatacggactgc | ccattttgtctacgggacga | 52 |
| *Hprt* | HPRT | tcctcctcagaccgctttt | aacctggttcatcatcgctaa | 95 |
| *Rn18S* | 18S rRNA | gccgctagaggtgaaattctt | cgtcttcgaacctccgact | 93 |
| *Tbp* | TBP | gggagaatcatggaccagaa | gatgggaattccaggagtca | 97 |
| *Agt* | AGT | cggaggcaaatctgaacaac | tcctcctctcctgctttgag | 84 |
| *Avp* | AVP | gctgccaggaggagaactac | aaaaccgtcgtggcactc | 88 |
| *Avpr1a* | V1aR | gggataccaatttcgtttgg | aagccagtaacgccgtgat | 31 |
| *Avpr1b* | V1bR | cattgtgctggcctacattg | tggtgaaagccacattggta | 104 |
| *Avpr2* | V2R | cctggtgtctaccacgtctg | gcaccagactggcatgtatct | 27 |
| *Crh* | CRH | gaggcatcctgagagaagtcc | atgttaggggcgctctcttc | 34 |
| *Crhr1* | CRHR1 | gggccattgggaaacttta | atcagcaggaccaggatca | 81 |
| *Cyp11b1* | CYP11B1 | gccatccaggctaactcaat | cattaccaaggggggttgatg | 11 |
| *Cyp11b2* | CYP11B2 | aagctcagacttggtgcttca | cggcccatggagtagagata | 3 |
| *Fkbp5* | FKBP5 | aaacgaaggagcaacggtaa | tcaaatgtccttccaccaca | 97 |
| *Hsd11b1* | 11βHSD1 | tctacaaatgaagagttcagaccag | gccccagtgacaatcacttt | 1 |
| *Hsd11b2* | 11βHSD2 | cactcgaggggacgtattgt | gcaggggtatggcatgtct | 26 |
| *Mc2r* | MC2R | caccacaatcctctaccctca | cctctccttggctttgtcac | 96 |
| *Nr3c1* | GR | gacgtgtggaagctgtaaagt | catttcttccagcacaaaggt | 56 |
| *Nr3c2* | MR | ttcggagaaagaactgtcctg | cccagcttctttgactttcg | 50 |
| *Pomc* | POMC | agtgccaggacctcacca | cagcgagaggtcgagtttg | 62 |
| *Serpina6* | CBG | ccaccaaagacactcccttg | ggtgtacaggagggccatt | 40 |
| *Srd5a1* | 5α-reductase1 | gggaaactggatacaaaataccc | ccacgagctccccaaaata | 41 |
| *StAR* | StAR | aaggctggaagaaggaaagc | ccacatctggcaccatctta | 2 |

***Abbreviations***: GAPDH; glyceraldehyde 3-phosphate dehydrogenase, HPRT; hypoxanthine-guanine phosphor-ribosyltransferase, rRNA; ribosomal RNA, TBP; TATA-binding protein, AGT; angiotensinogen, AVP; vasopressin, V1aR; vasopressin receptor 1A, V1bR; vasopressin receptor 1B, V2R; vasopressin receptor 2, CRH; corticosterone releasing hormone, CRHR1; CRH receptor 1, CYP11B1/2; cytochrome P450, family 11 subfamily b polypeptide 1/2, FKBP5; FK506 binding protein 51, 11βHSD1/2; *11β*-Hydroxysteroid dehydrogenase type 1/2, MC2R; melanocortin receptor 2, GR; glucocorticoid receptor, MR; mineralocorticoid receptor, POMC; pro-opiomelanocortin, CBG; corticosteroid binding globulin, StAR; steroidogenic acute regulatory protein.

**Figure S1.** A) Adrenal gland weight and (B) adrenal gland mRNA abundance for *Cyp11b1* from mice fed either a 0.3% Na diet (Control, open circles) or a 3% Na diet (High Salt; grey circles) for 14 days. Individual values are shown with group mean±SD; statistical comparisons were made using Student's unpaired *t* test with two-tailed p values stated.

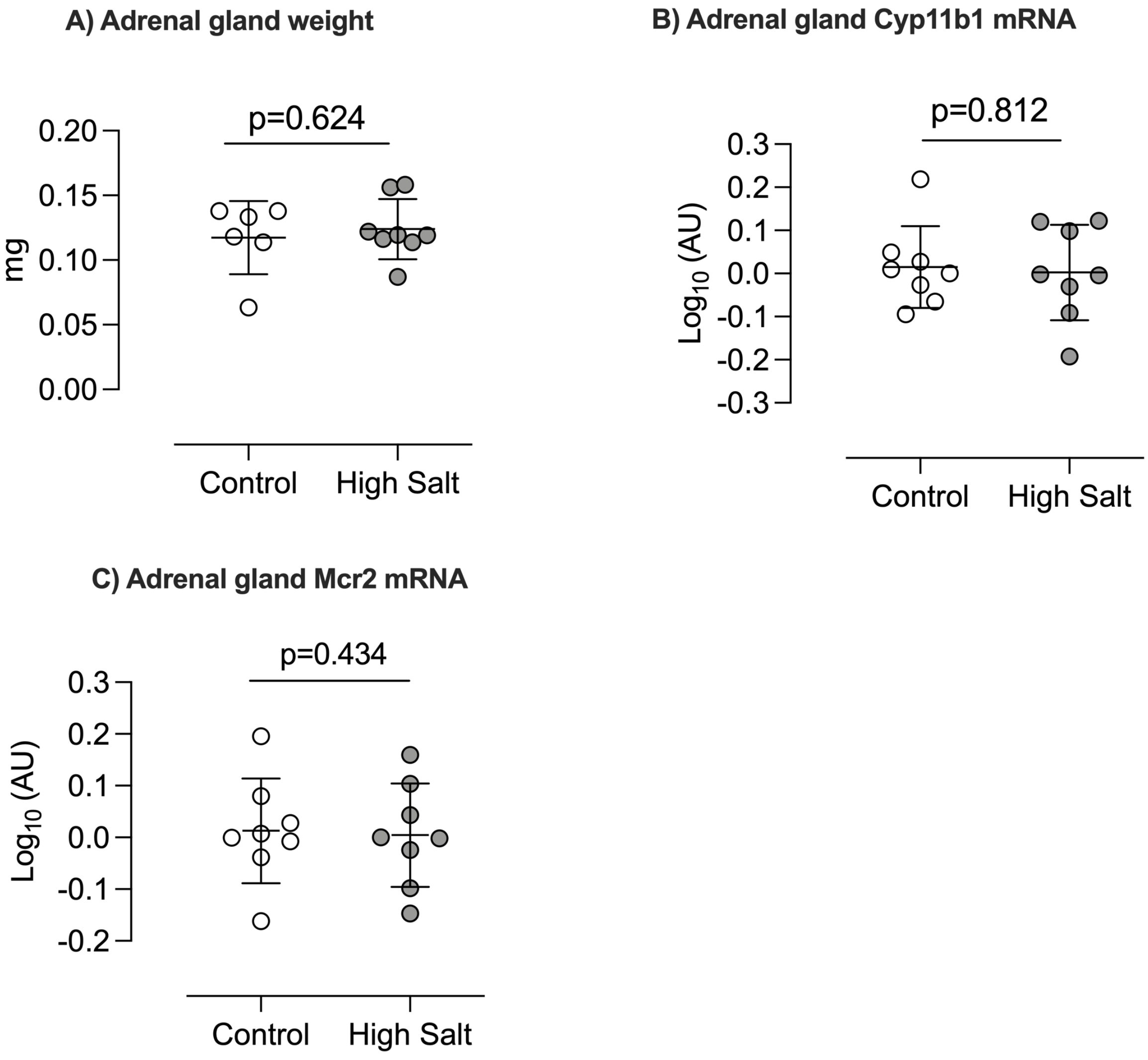

**Figure S2.** A) plasma aldosterone and mRNA abundance for B) hepatic *Agt;* C) adrenal gland *Cyp11b2; D)* kidney cortex *Nr3c2 and E)* kidney medulla *Nr3c2* in mice fed either a 0.3% Na diet (Control, open circles) or a 3% Na diet (High Salt; grey circles) for 14 days. Individual values are shown with group mean±SD; statistical comparisons were made using Student's unpaired *t* test with two-tailed p values stated.

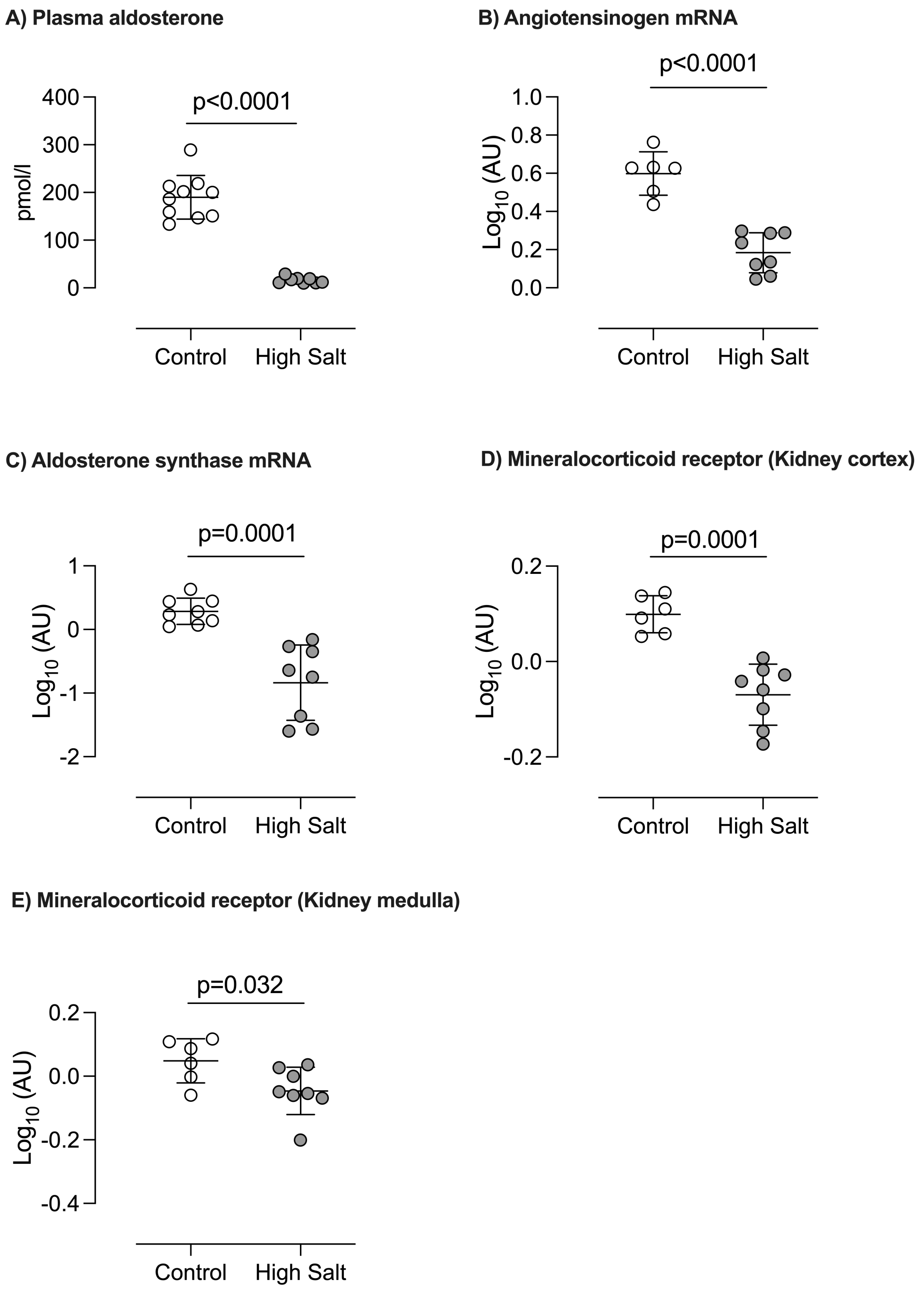

**Figure S3.** A ) Plasma CBG binding capacity; B) liver mRNA abundance for *SerpinA6* and from adult male mice fed either 0.3% Na diet (Control, open circles) or 3% Na diet (High Salt; grey circles) for 2 weeks. Individual values are shown with group mean±SD; statistical comparisons were made using Student's unpaired *t* test with two-tailed p values stated.

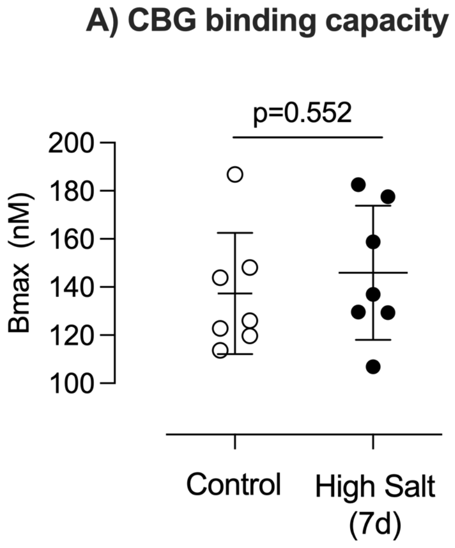


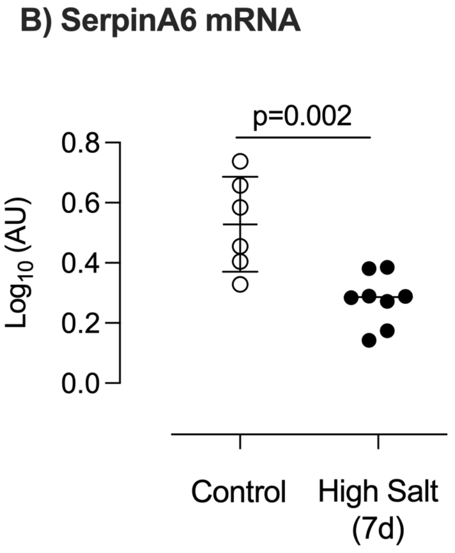

**Figure S4.** mRNA abundance in kidney cortex and kidney medulla of A) *Nr3c1* (glucocorticoid receptor GR); B) *Hsd11b1* (11β hydroxysteroid dehydrogenase type 1) and C) *Fkbp5* (FK506 binding protein 5) from mice fed either 0.3% Na diet (Control, open circles) or 3% Na diet (High Salt; grey circles) for 2 weeks. Individual values are shown with group mean±SD; statistical comparisons were made using Student's unpaired *t* test with two-tailed p values stated.

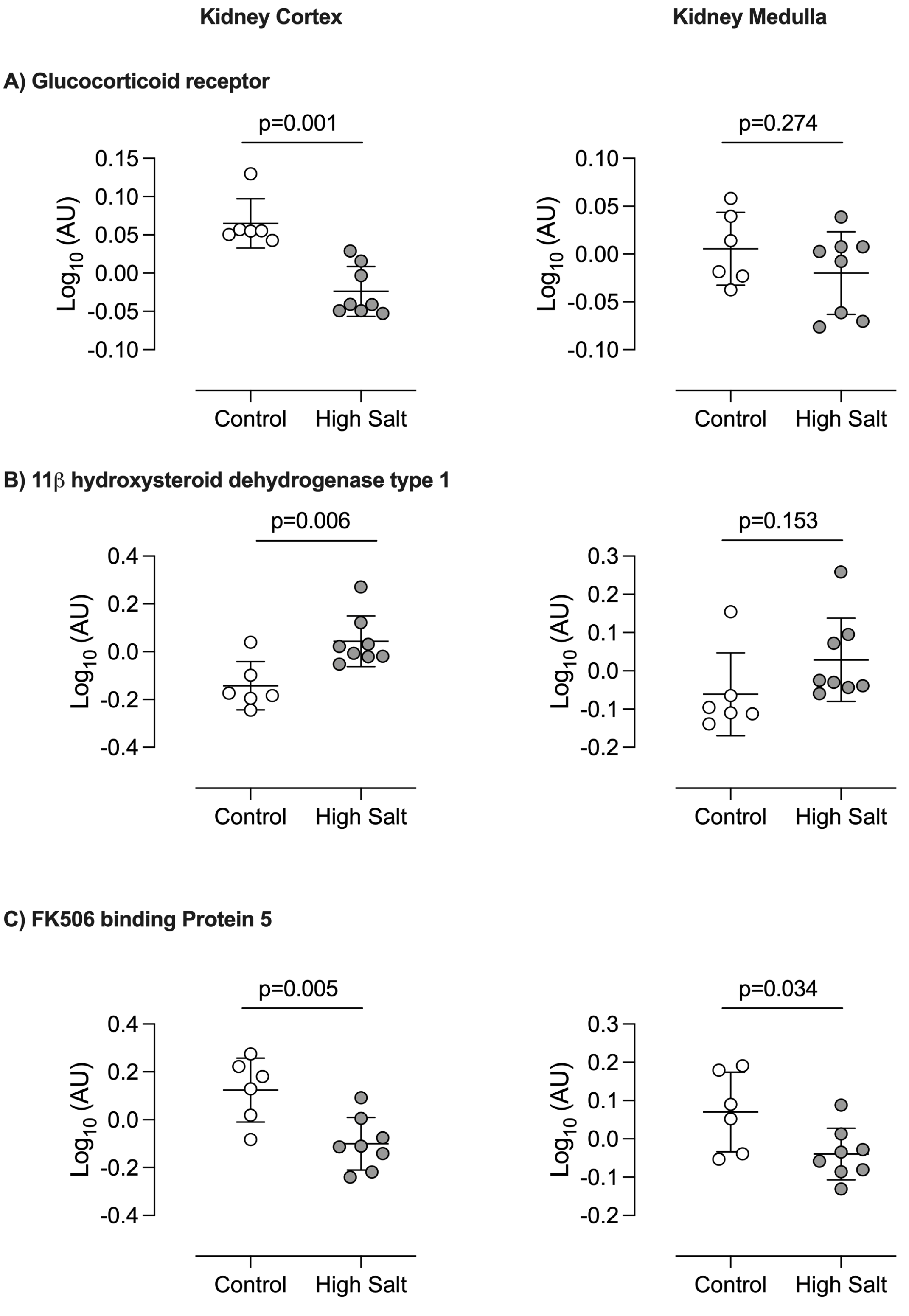

**Figure S5.** mRNA abundance in heart and aorta of A) *Nr3c1* (glucocorticoid receptor GR); B) *Hsd11b1* (11β hydroxysteroid dehydrogenase type 1) and C) *Fkbp5* (FK506 binding protein 5) from mice fed either 0.3% Na diet (Control, open circles) or 3% Na diet (High Salt; grey circles) for 2 weeks. Individual values are shown with group mean±SD; statistical comparisons were made using Student's unpaired *t* test with two-tailed p values stated.

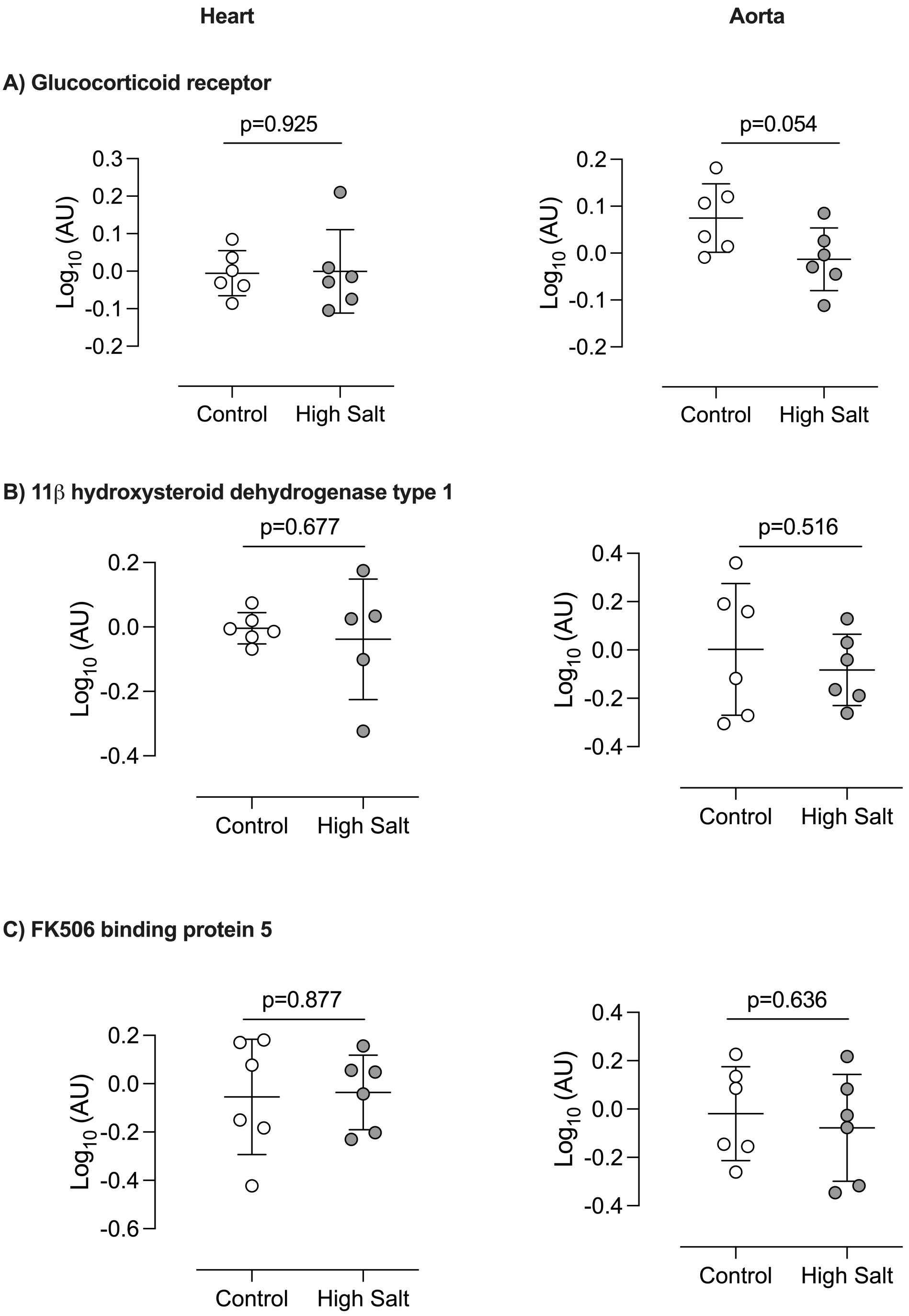

**Figure S6.** mRNA abundance in adrenal gland and hypothalamus of A) *Nr3c1* (glucocorticoid receptor GR) and B) *Fkbp5* (FK506 binding protein 5) from mice fed either 0.3% Na diet (Control, open circles) or 3% Na diet (High Salt; grey circles) for 2 weeks. Individual values are shown with group mean±SD; statistical comparisons were made using Student's unpaired *t* test with two-tailed p values stated.

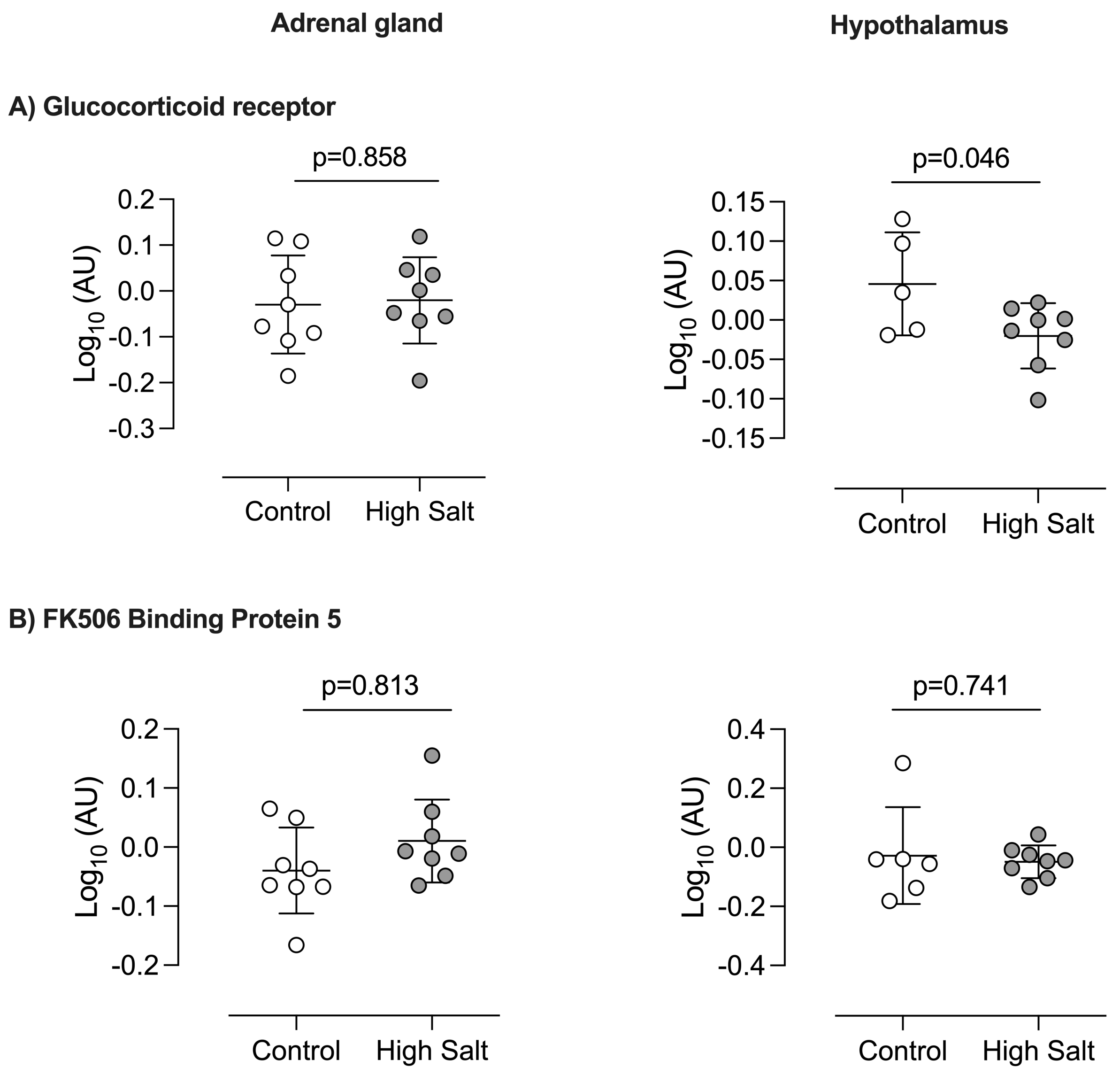

**Figure S7.** mRNA abundance in omental white adipose tissue of A) *Nr3c1* (glucocorticoid receptor GR) and B) *Fkbp5* (FK506 binding protein 5) from mice fed either 0.3% Na diet (Control, open circles) or 3% Na diet (High Salt; grey circles). Individual values are shown with group mean±SD; statistical comparisons were made using Student's unpaired *t* test with two-tailed p values stated.

**A) Glucocorticoid receptor**

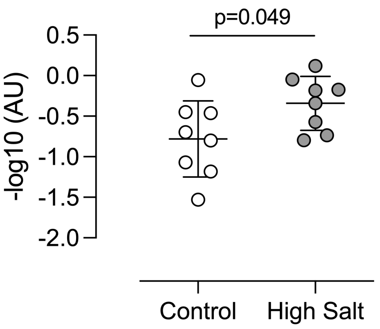


**B) 11β hydroxysteroid dehydrogenase type 1**

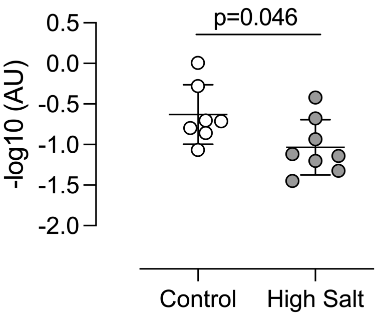


**C) FK506 binding protein 5**

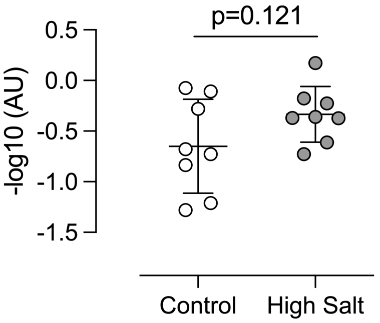
